# Corticohippocampal Neuroenergetics and histomorphology in aluminium-induced neurotoxicity: Putative therapeutic roles of ascorbic acid and nicotine

**DOI:** 10.1101/2020.07.09.195495

**Authors:** Gbadamosi Ismail Temitayo, Omotoso Gabriel Olaiya

**Affiliations:** Division of Neurobiology, Department of Anatomy, Faculty of Basic Medical Sciences, College of Health Sciences, University of Ilorin

**Keywords:** Alzheimer, Anxiety Neuroenergetics, Nicotine, Ascorbic acids

## Abstract

Alzheimer’s disease (AD) is the most common cause of dementia and is hallmarked by β-amyloid plaque and neurofibrillary tangles deposition in the central nervous system. The complex mechanism that underlies AD pathogenesis has made the development of a definitive cure futile. Exploring the possible therapeutic advantages of combining two neuromodulatory molecules with different mechanisms of neuroprotection is an interesting way of drug discovery. Ascorbic acid (AA), a potent antioxidant molecule, and nicotine (NIC), an allosteric modulator of nAChRs, have both been documented to independently proffer neuroprotection in experimental and clinical neurodegenerative cases. This study elucidated the putative therapeutic advantages of combining ascorbic acid and nicotine as a treatment regimen against the aluminium-induced Alzheimer-like corticohippocampal histopathology, anxiety, and perturbed neuroenergetics in rats induced with

Rats treated with 100 mg/kg aluminium chloride for 28 days presented with significantly increased stretch attend posture frequency and centre square entry. Aluminium significantly depleted the activities of glucose-6-phosphate dehydrogenase (G6PDH) while increasing lactate levels. Corticohippocampal histomorphology of these animals showed poor histoarchitecture, increased congophilic and argentophilic densities that were coupled with increased anti-NSE immunopositivity. Animals post-treated with NIC (10mg/kg) and AA (100mg/kg) for 28 days presented with reduced anxiety level and improved corticohippocampal histomorphology. AA normalized G6PDH and lactate levels while the congophilic density was reduced by NIC. Corticohippocampal argentophilic density anti-NSE immunopositivity were also normalized by AA+NIC.

The findings from this study have shown that a combination of ascorbic acid and nicotine effectively mitigated aluminium-induced corticohippocampal histopathology and perturbed neuroenergetics.

## Introduction

Alzheimer’s disease (AD), the most common cause of dementia (Elahi and Miller, 2017), is a progressive and debilitating type of neurodegenerative disorder (NDD), classically characterized by β-amyloid plaque and neurofibrillary tangles deposition in the brain (Weller and Budson, 2018). The most devastating aspect of this disease is the manifestation of impaired behavioural phenotypes, like memory loss, anxiety and reduced cognition (Mu and Gage, 2011), that often result in diminished life quality (Chalazonitis and Rao, 2018). Even worse, it has no known cure, and about a decade of life span after being diagnosed (Pepeu and Giovannini, 2010). Consequently, the ultimate goal of finding a drug therapy against AD is to prevent the development or significantly delay the onset (or progression) of the disease pathology, while enhancing or averting the deterioration in cognitive and memory functions in people diagnosed with the disease.

The observed behavioural manifestation in AD patients can be attributed to the neurochemical imbalances (Gsell et al., 2005) that culminate in compromised histomorphology and functional integrity of neurons and surrounding glial cells in vital brain areas like the prefrontal cortex (PFC) and hippocampus (HIP) (Bazzigaluppi et al., 2018; Pang et al., 2017). Sustained neural oxidative stress, for example, activates beta-secretase (BACE) activities (Jo et al., 2010; Tamagno et al., 2008) and inhibits cytochrome oxidase (Kann, 2016), thereby shifting amyloid precursor protein (APP) processing to an amyloidogenic derivative (Pohanka, 2018). Oxidative stress also induces tau hyperphosphorylation, which destabilizes microtubules by decreasing tau binding affinity, resulting in the formation of NFTs (Liu et al., 2015). Other neurochemical events that have been elucidated in the pathogenesis of AD include nitrosative stress (Serini and Calviello, 2015), dysfunctional cholinergic transmission (Posadas et al., 2013), exacerbated lipid peroxidation (Sultana et al., 2013), neuroinflammation (Heneka et al., 2010), and perturbed iron homeostasis (Lane et al., 2018), to mention a few. Furthermore, in AD’s pathophysiology, neuronal and glial cells tend to use the energetically inefficient form of burning glucose through glycolysis as a “makeshift” response to increase metabolic processes (An et al., 2018; Atlante et al., 2017). The shift in energy metabolism becomes a disadvantage since the loss of the adaptive advantage afforded by elevated aerobic glycolysis exacerbates the pathophysiological processes, such as lactate accumulation (Ross et al., 2010), associated with AD, making the brain susceptible to Aβ-induced neurotoxicity and leading to cell death and dementia. The aforementioned neurochemical perturbations culminate in the histomorphological changes observed in AD brain.

The complex mechanism that underlies AD pathogenesis has made the development of a definitive cure futile. However, a few medical treatments have been approved for AD and these act to control symptoms rather than alter the course of the disease (Briggs et al., 2016). Much of AD research over the past decade has concentrated on disease-modifying treatment that will change the course of the disease rather than on symptoms alone, but the lack of effective disease-modifying medications from these trials illustrates the challenges of developing a therapeutic agent with the ability to alter the course of a disorder as nuanced as AD. There are several molecular approaches to the management of AD. Exploring the possible therapeutic advantages of combining two neuromodulatory molecules with different mechanisms of neuroprotection is an interesting way of drug discovery (Ho and Chang, 2004; Zaky et al., 2017). Ascorbic acid (AA), a potent antioxidant molecule (Arrigoni and De Tullio, 2002; Dorota Majewska and Bell, 1990; Foyer, 2017), and NIC (NIC), an allosteric modulator of nAChRs (Albuquerque et al., 2009; De Biasi and Dani, 2011; Mansvelder and McGehee, 2002), have both been documented to independently proffer neuroprotection in experimental and clinical neurodegenerative cases (Alkadhi, 2011; Ho and Chang, 2004; Olajide et al., 2017; Ryan et al., 2001).

NIC is able to modulate cholinergic transmission and reduce memory decline in an experimental AD model by binding with nAChRs and upregulation of ChAT (Hernandez and Terry, 2005; Wonnacott, 1990). Furthermore, NIC has been reported to breakdown preformed amyloid plaques by inhibiting the activities of BACE-1 (Srivareerat et al., 2011). However, despite the broad range of the therapeutic advantages of NIC, it has been labelled as being neurotoxic (Ferrea and Winterer, 2009). In addition to some of our previous reports, there are several lines of evidence that NIC promotes the production of reactive oxygen and nitrogen species, promotes lipid peroxidation and ultimately compromises the neural antioxidant defence system (Carlson et al., 2001; Elsonbaty and Ismail, 2020; Gbadamosi et al., 2016; Omotoso et al., 2018). Hence, NIC tendency to drive oxidative stress undermines its therapeutic advantages. AA is a potent antioxidant vitamin (Covarrubias-Pinto et al., 2015) whose deficiency had been linked to several neurodegenerative diseases (Olajide et al., 2017b; Parle and Dhingra, 2003). Owing to its antioxidant properties, dietary supplementation of AA has been associated with reduced prevalence and incidence of AD. Several studies have noted the neuroprotective role of AA in different models of AD. However, AA enhances the uptake of iron (Lane and Richardson, 2014).

One of the many approaches used in drug discovery is to induce Alzheimer-like neuropathologies in experimental animals using different models. Aluminium chloride is a chemical substance that has been reported to exhibit Alzheimer-like phenotypes following acute and chronic toxicity (Shati et al., 2011; Singh and Goel, 2015; Taïr et al., 2016). Aluminium has been reported to promote reactive oxygen species (ROS) production (Igbokwe et al., 2020a), intensify lipid peroxidation (Sumathi et al., 2015), disrupt cholinergic neurotransmission (Ghorbel et al., 2016), trigger neuroinflammation (Cao et al., 2016), promote chromatolysis (Elizabeth et al., 2020), stimulate glial activation (Akinrinade et al., 2015), promote neuronal cell death, and cause anxiety, cognitive decline and memory loss. In this study, we explored the changes in anxiety levels, corticohippocampal histomorphology and neuroenergetics following aluminium-induced neurotoxicity, and then elucidated the putative therapeutic advantages of combining ascorbic acid and nicotine as a treatment regimen against the induced neuropathology.

## Methodology

### Ethical statement

Following the ethical approval given by the University of Ilorin Ethical Review Committee (UERC) with reference number UERC / ASN/2018/1473, this investigation was conducted at the Department of Anatomy. Animal handling procedure for the care and use of laboratory animals was enforced in full adherence with the National Institutes of Health Handbook (NIH Guidelines No.2.4.3.8023, updated 1978).

### Chemicals

AA (Cat.No.:1043003) and AlCl3 salt (Cat. No.: 563919) were obtained from Sigma - Aldrich (USA). 95% Nicotine (Cat. No.:412634) was purchased from British Drug House (BDH) Chemical Ltd, London. Distilled water from Lab Trade Limited (LTL), Ilorin Nigeria was used as a medium for the degradation and administration of nicotine, ascorbic acid and AlCl3 to experimental animals.

### Experimental animals

For the present study, forty (40) Male Wistar rats, obtained from the Faculty of Veterinary Sciences, University of Ilorin, were used. The animals were acclimatized and housed under standard laboratory conditions with liberal access to standard rat feed and drinking water at the Animal Holding Facility of the Faculty of Basic Medical Sciences, University of Ilorin.

### Animal treatment

The animals were split into five (A-E) groups of eight animals each. Group A (control) was treated with distilled water daily for 4 weeks. Aluminium toxicity was achieved in groups B-E through a daily oral infusion of 100mg/kg of AlCl3 for four weeks. Groups C (AA), D (NIC) and E (AA+NIC) were then post-treated with AA (100mg/kg daily), NIC (10mg/kg daily) and a combination of NIC (10mg/kg daily) and AA (100mg/kg daily), for four weeks. AlCl3 and AA were given orally using oral gavage while NIC was intraperitoneally administered using insulin syringes. Group E received AA six hours after being treated with NIC. At the end of the fourth week, groups A and B were behaviorally tested and sacrificed, while groups C-E were tested and sacrificed at the end of the eighth week (figure 1a).

**Fig 1.**
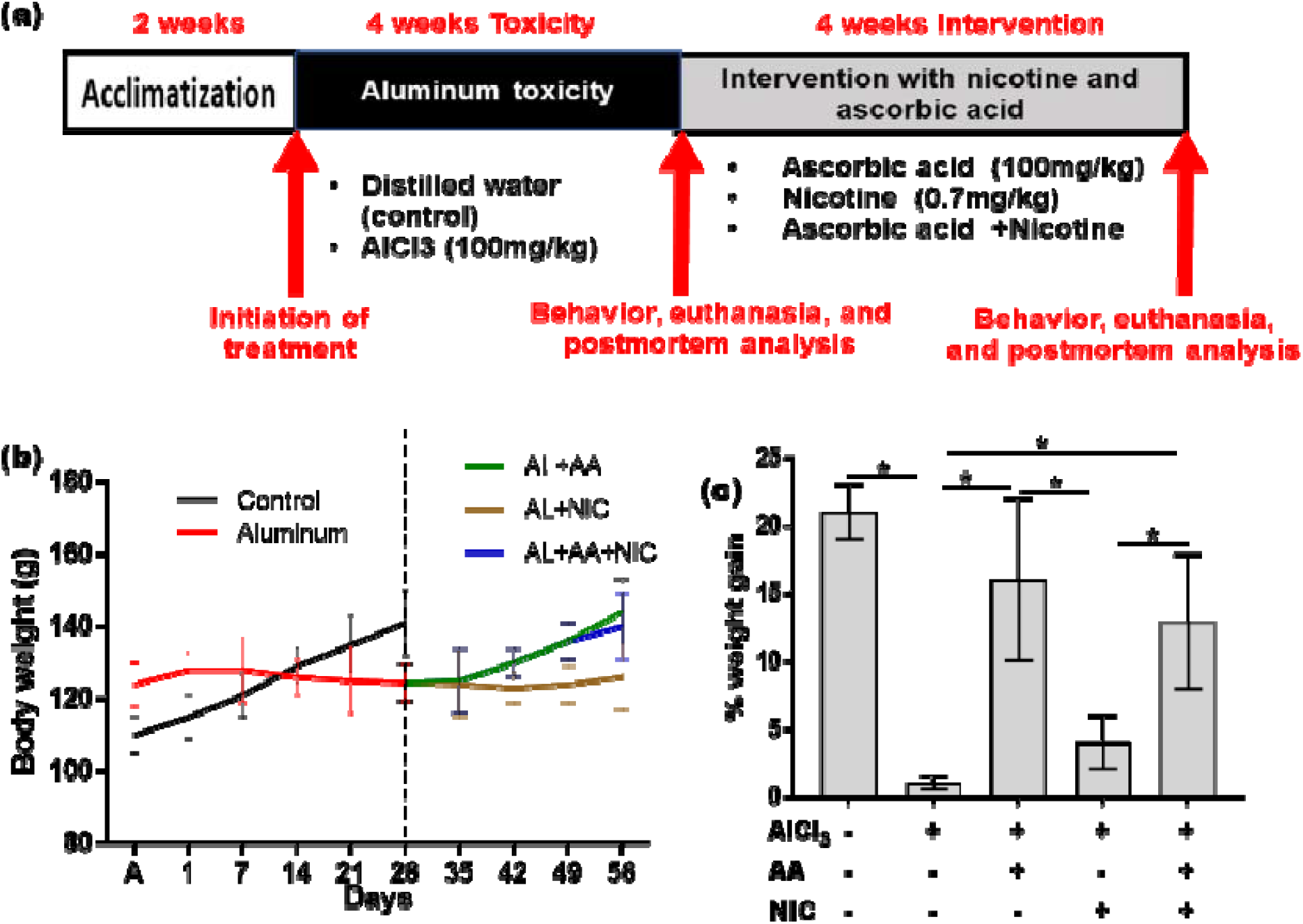
Experimental design (a), rate of weight change (b) and relative weight gain (c) of experimental Wistar rats induced with aluminium toxicity and then treated ascorbic acid, nicotine, or a combination of both. a) animals were acclimatized for two weeks and then orally induced with aluminium toxicity for four weeks. The animals were then treated with AA, NIC, and AA+NIC. b) during the first four weeks, the animals treated with aluminium presented with a linear decrease in body weight. Treatment with either of AA or A+NIC, during the following four weeks, caused a linear increase in body weight. c) Aluminium and NIC groups presented with significantly lower body weight relative to the control, AA and AA+NIC group. * is a significant value of p<0.05.

### Open Field Test

The open-field test (OFT) was used to quantify the exploratory drive and anxiety levels of the experimental animals (Gould et al., 2009). The OFT apparatus consists of an open-top box measuring 100cm in length and width, and 50cm in height. The apparatus floor was marked out into 16 equal squares with a black line and one centre square with a red line. During the test, each animal was placed in the centre square and allowed to explore for 10 minutes while a camera suspended from the ceiling captured the activities of the animals. The floor of the open field paradigm was wiped clean with 70% ethanol before and after each animal. The video was later analyzed to obtain data which included the number of lines crossed, centre square entry, centre square duration, rearing frequency, rearing duration, and stretched attend posture frequency

### Animal sacrifice and tissue processing

At the end of the behavioural tests, the animals in each group were sacrificed in two categories. Rats used for histological, histochemical and immunohistochemical studies (3 rats per group) were euthanized by intramuscular injection of ketamine. The animals’ thoracic cavities were opened to expose the heart for transcardial perfusion. The perfusion was carried out through the left ventricle, starting with normal saline to remove blood, followed by 4% paraformaldehyde (PFA). The animals were then decapitated, and the brain tissues were dissected out of the skull. Excised brains were then rinsed in 0.25 M sucrose three times each and then post-fixed in 4% PFA for 24 hours. The prefrontal cortex (PFC) and hippocampus were dissected out and were preserved in 20% sucrose (in PBS) containing 0.02% sodium azide at 4°C until further processing. PFA-fixed tissues were dehydrated, cleared, infiltrated and embedded in paraffin wax.

### Histology and histochemical procedure

The corticohippocampal general histomorphology, amyloid plaques and axodendritic morphology were demonstrated using H & E (hematoxylin and eosin) stain, Congo red stain and Bielschowsky silver stain, respectively. Paraffin-embedded tissues were sectioned at 5 microns, floated, picked with glass slides, dried, deparaffinized in two changes of xylene, rehydrated through decreasing grades of alcohol (100%, 95%, 70%, and 50%) and rinsed in distilled water. Sections for H&E stain were stained in Mayer hematoxylin, rinsed in running tap water, briefly blued in 95% alcohol, rinsed with distilled water, counterstained in 025% Eosin Y, rinsed severally, dehydrated, cleared, and mounted in DPX. Sections for Congo Red stain were stained in 0.5% Congo red in 50% ethanol solution, rinsed, counterstained in Mayer’s hematoxylin, rinsed again, dehydrated, cleared, and mounted in DPX. Slides for Bielschowsky silver stain were incubated in ammoniacal silver solution (6% ammonium hydroxide in 20% silver nitrate), then passed through 1.5% ammonium hydroxide, developed in treated ammoniacal solution, dipped in 1.5% ammonium hydroxide, rinsed in distilled water, put in 5% sodium thiosulphate to stop the silver reaction, rinsed, dehydrated, cleared, and mounted.

### Immunohistochemical procedures

PFC and Hippocampal sections were taken from paraffin blocks to charged glass slides. Heat-induced antigen retrieval was performed using citrate buffer solution (10nM citric acid, 0.05% Tween 20, pH 6.0) at 98° C, to reveal masked epitope. Subsequently, non-specific protein reaction blocking was performed in 10% normal goat serum in 10 mM PBS + 0.03% Triton-X 100 and 1% bovine serum albumin (BSA) for 2 hours at room temperature. The endogenous peroxidase block was done using 0.3% hydrogen peroxide in TBS (15 min). Sections were then incubated in 500 μl of rabbit anti-NSE solutions (1:100 dilution each), diluted in blocking buffer (10% goat serum with 1% BSA and 0.1% Triton X-100 in 10 mM PBS) overnight at 4 °C. After adequate washes, appropriate goat anti-rabbit HRP conjugated secondary antibodies were diluted in TBS + 1% BSA was applied to slides for an incubation period of 1 hour at room temperature. All incubations following antigen retrieval were done in a humidity chamber to avoid drying of sections. The immunogenic reaction was developed using 3’3’ Diaminobenzidine tetrachloride (DAB). Sections were then counterstained in hematoxylin, washed, dehydrated in absolute alcohol, cleared in xylene and mounted in DPX.

### Colourimetric assay for biochemical studies

Rats for enzymatic studies (5 rats per group) were sacrificed by cervical dislocation to prevent an anaesthetic agent from meddling with biochemical redox. Following careful decapitation, the skulls were dissected to remove the brains. The brain tissues were rinsed in 0.25M sucrose and the dissected on a cold plate to obtain the PFC and hippocampi following which they were stored in 0.25M sucrose solution at 4° C. The brain tissues were homogenized in 0.25 M sucrose using an automated homogenizer at 4° C. Homogenates were centrifuged at 2795g for 10 min at 4° C. The supernatants were aspirated to quantify enzyme activities. Activities of glucose-6-phosphatase dehydrogenase (G6PDH) and lactate were measured by immunosorbent quantification using the spectrophotometric technique as specified in their respective assay kits.

### Light Microscopy, Densitometric and Statistical Analysis

Histological, histochemical and immunohistochemical analyses of the hippocampus and PFC were captured using Olympus binocular research microscope (Olympus, New Jersey, USA) which was connected to a 5.0 MP Amscope camera (Amscope Inc, USA). Densitometric analysis to quantify congophilic, argentophilic and NSE staining intensities were determined from captured PFC and hippocampal images using the Image J (NIH, USA). For each section, the staining intensity was determined by the mean grey area (in image J) around the external pyramidal layer and CA3 (using cellular morphology and layer-dependent cell densities) for five different fields of view at high-power magnification. All quantitative data were analyzed using GraphPad Prism software (Version 7). ELISA and staining intensity outcomes were plotted using one-way ANOVA with Tukey’s multiple comparisons test. Significance was set at p < 0.05*. The results were presented in graphs with error bars to show the means and standard error of means, respectively.

## Results

### AA+NIC post-treatment improved body weight

The bodyweight of the Wistar rats in this study was obtained and recorded at a weekly interval. Analysis of the bodyweights showed that aluminium induced a weight loss. The line graph revealed that the weight of animals in the control group increase linear fashion through the four weeks of treatment (fig. 1b). The aluminium treated rats presented with an increase in body weight during the first week. The body weight was maintained at a constant level for another week. The bodyweight began to reduce in the third week of treatment. The body weight was maintained at a fairly constant level during the fourth week (fig. 1b). Analysis off the percentage increase in boy weight showed that aluminium had a significantly lower increase in body weight when compared to the control group (p<0.05). The finding suggests that aluminium treatment has a weight loss effect on the experimental animals (fig. 1c). Animals treated with ascorbic acid showed a continuous linear increase in body weight for four weeks. This resulted in a significant increase in body weight when compared to the aluminium group (p<0.05). Nicotine, however, did not have a significant effect on the bodyweight following aluminium toxicity. Interestingly, the animal group that was treated with a combination of AA and NIC presented with a continuous linear increase in body weight over the four weeks of treatment. This surmounted to a significant body weight increase compared to the aluminium group (p<0.05). Comparatively, the AA+NIC groups had significantly higher percentage weight gain when compared to the NIC treated group (fig. 1c). This finding suggests that AA played a role in sustaining the increase in body weight of the animals in the presence or absence of NIC treatment.

### AA+NIC treatment mitigated aluminium-induce anxiety

Following the completion of the drug treatment, the animals were subject to the open field behavioural paradigm. Aluminium treated rats present with no significant changes in centre square entry and duration, as well as rearing frequency and duration. One may think that aluminium treatment for four weeks at the dose adopted in this study has no significant effect on the exploratory drive of the animals. However, the animals in the aluminium group presented with significantly higher stretch attend posture frequency, suggesting anxiety behavioural. This finding suggests that aluminium was anxiogenic, thereby causing the insignificant changes in exploratory drive observed in the aforementioned behavioural indices in the open field. Test. Treatment of these animals with ascorbic acid nicotine treatment regimen showed some anxiolytic manifestations.in the open field test, AA, NIC, and AA+NIC presented with a higher number of lines crossed and stretch-attend posture frequency. However, the observed increase was not significant. Comparatively, there was no significant difference in the centre square entry and duration, as well as the rearing frequency and duration. Interestingly, AA and NIC treatment, individually or in combination, mediated a notable reduction in stretch attend posture frequency. This decrease was significant in the AA+NIC group when compared to the aluminium group (figure2)

**Fig 2.**
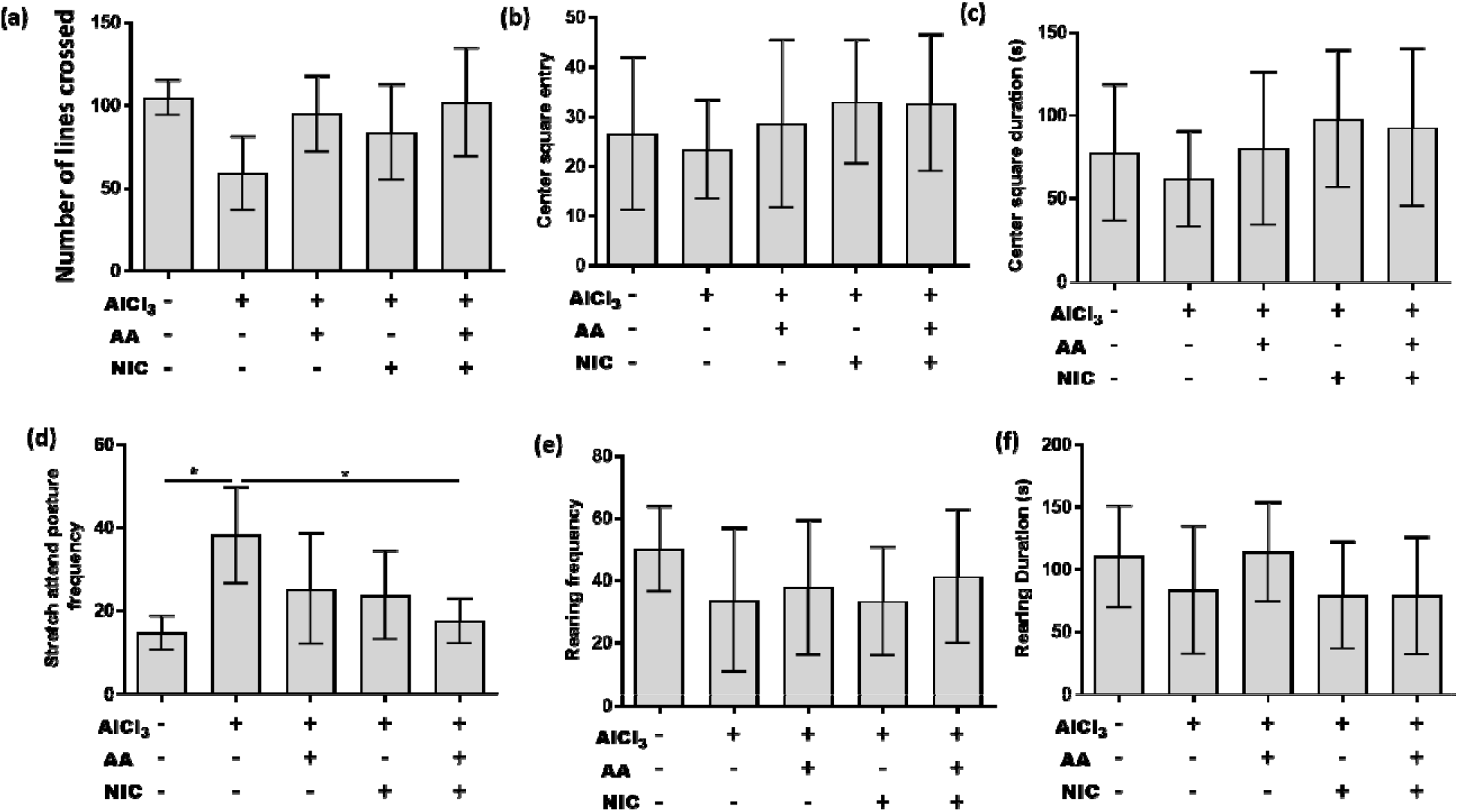
Behavioral manifestation of animals in the open field behavioural paradigm. The aluminium only group presented with a lower number of lines crossed (a), centre square entry (b), centre square duration (c), rearing frequency (e), and rearing duration (f), relative to the control group and treated group. Stretch attend posture frequency (d) was significantly higher in the aluminium group when compared to the control group and AA+NIC group. * is significant value of p<0.05).

### AA+NIC restores perturbed corticohippocampal neuroenergetics

A consistent supply of molecular oxygen tissues at a subcellular level is crucial to normal physiological functioning. In the pentose phosphate pathway, glucose-6-phosphate dehydrogenase performs an essential function in making reduced energy available. Findings from this showed that when compared to the control group, the aluminium group had a significantly reduced G-6-PDH (p<0.05 in both brain regions). This finding suggests that aluminium perturbed energy metabolism from oxidative phosphorylation. The perturbation may be the result of increasing energy demand following the toxicity mediated by aluminium. The glycolytic pathway, even though it does not produce much ATP, is quick source energy at a subcellular level. Hence, the level of lactate was assayed in the cortical and hippocampal lysate. The results showed that when compared to the control group, the aluminium group had a significantly higher level of lactate in the PFC (p<o.05) and hippocampus (p<0.05). This finding confirms the speculation that aluminium induced a paradigm shift in energy metabolism from oxidative phosphorylation to glycolysis in a bid to cope with the excessive energy demand induced aluminium toxicity. Although NIC played no significant role in modulating G-6-PDH and lactate activities, AA significantly reversed the depleted G-6-PDH activities and attenuated exacerbated lactate levels. When compared to the AL group, the AA group presented with a significant increase in G-6-PDH (p<0.05 in both brain regions). Similarly, the AA+NIC group showed a significant increase in G-6-PDH relative to the AL group (p<0.05 in both brain regions). In a like manner, AA significantly overturned the exacerbated lactate activities. Relative to the AL group, the AA group presented with a significantly reduced lactate (p<0.05 in both brain regions). Likewise, the AA+NIC group showed a significant reduction in LDH relative to the AL group (p<0.05 in both brain regions) (figure 3). These findings suggest that the ascorbic acid component of the AA+NIC treatment regimen enhanced energy derivation through oxidative phosphorylation, possibly through modulation G-6PDH activities in the pentose phosphate pathway.

**Fig 3.**
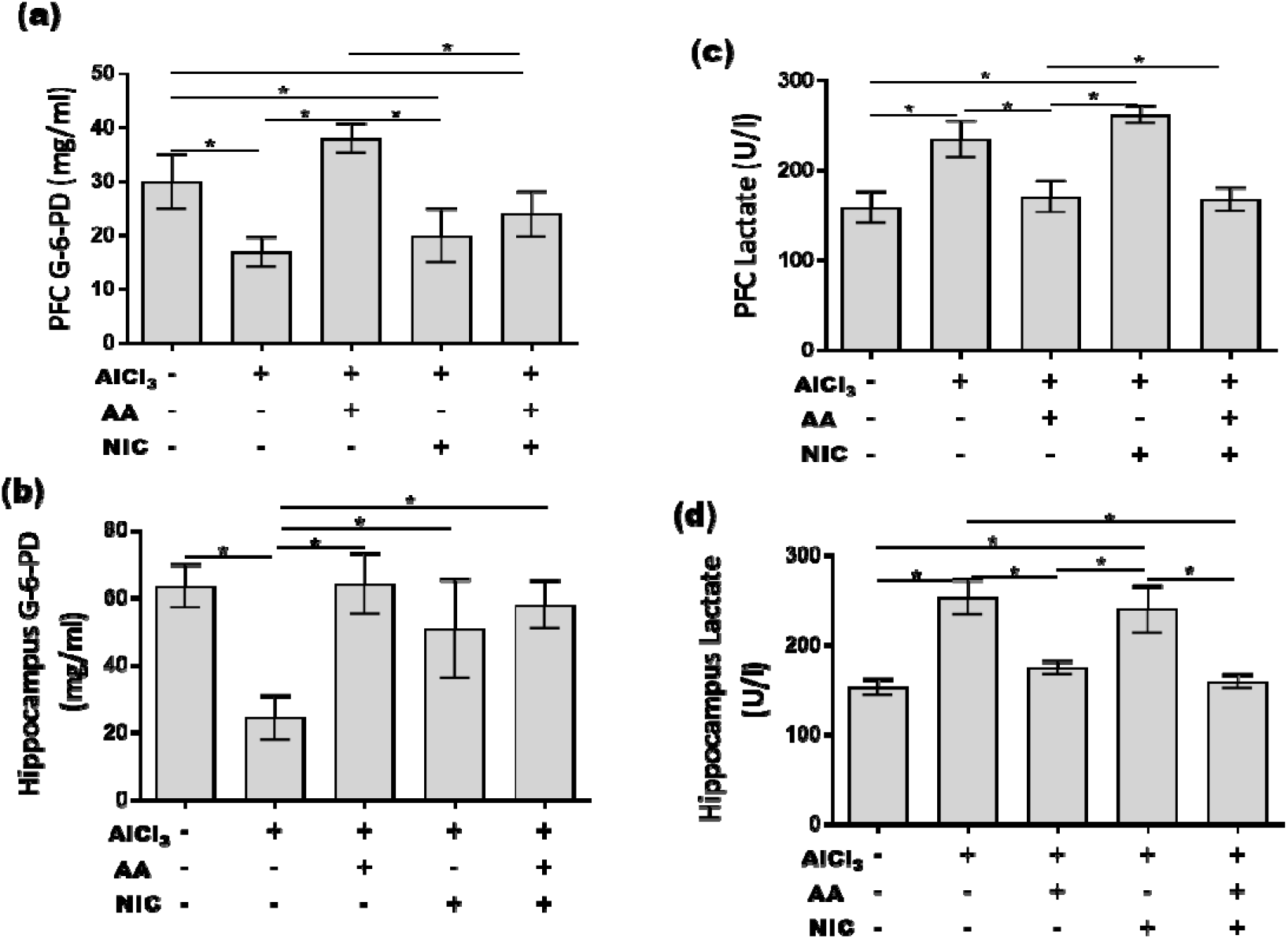
corticohippocampal energy metabolism in experimental animals induced with aluminium toxicity and then treated ascorbic acid, nicotine, or a combination of both. The aluminium group presented with a significantly reduced level of G-6-PDH when compared to the control and AA group in both PFC and hippocampus. Conversely, lactate level was significantly increased in the aluminium group when compared to the control, AA, and AA+NIC in both PFC and hippocampus.

**Fig 4.**
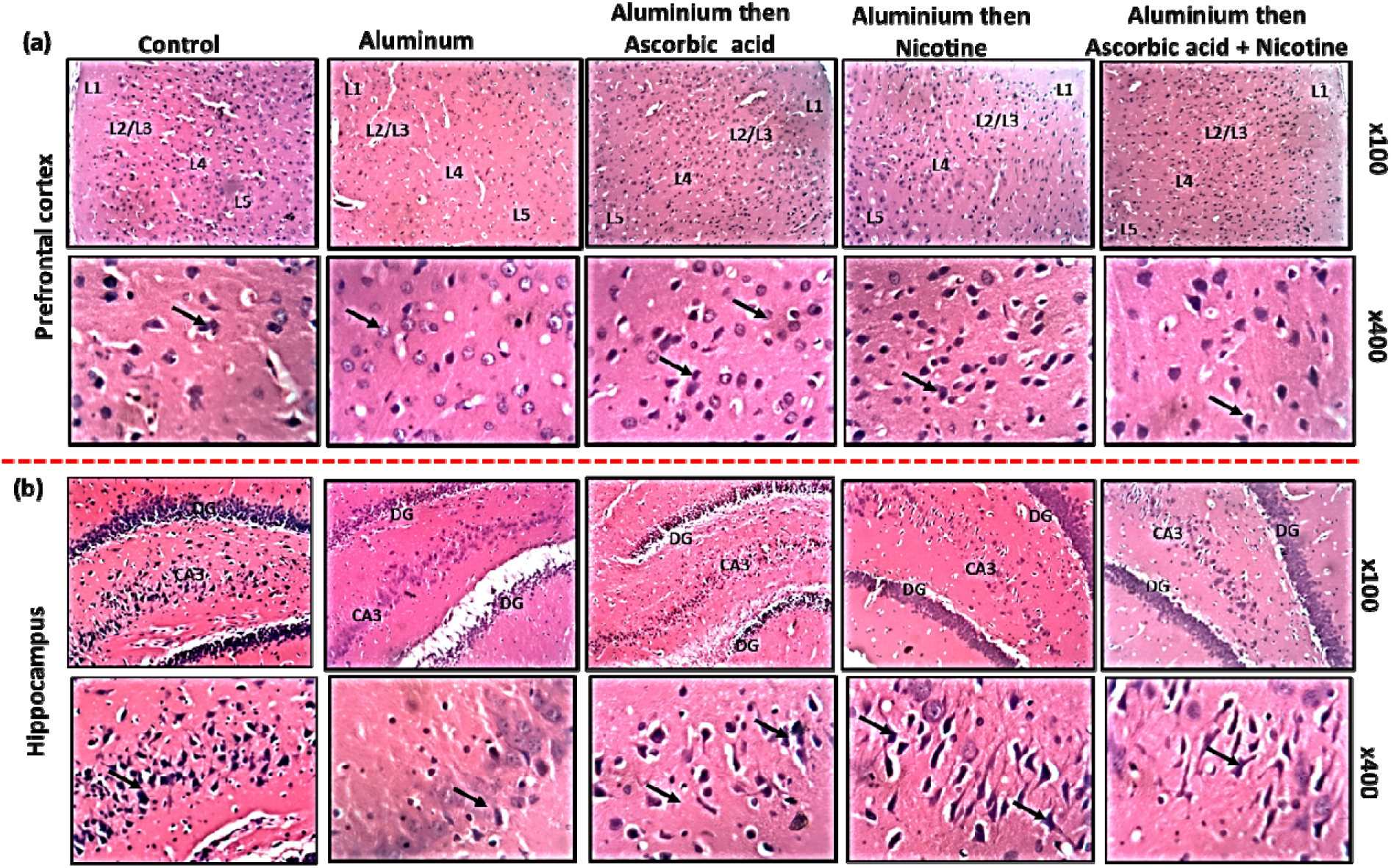
Representative photomicrograph of the general histology of the PFC (a) and hippocampus (b) of Wistar rats induced with aluminium toxicity and then treated ascorbic acid, nicotine, or a combination of both. a) The x100 magnification of the PFC shows a panoramic view of the cortical layers showing the molecular layer (L1), external granular and pyramidal layer (L2/L3), internal granular layer (L4) and internal pyramidal layer (L5). The x400 magnification reveals the external pyramidal layer with ubiquitously distributed pyramidal neurons (black arrows) with large soma and conspicuous apical and basal projections, interspersed with few granular neurons. The control group present with typical cortical delineation, cellularity and intact and well-characterized external pyramidal layer. The aluminium group presents with poor delineation and reduced cellularity across the cortical layers. The external pyramidal layer presents with neuropil fragments and reduced staining intensity. AA, NIC and AA+NIC posttreatment groups present with improved PFC histomorphology. b) The panoramic view (x100) of the hippocampus reveals the CA3 in the surrounding dentate gyrus (CA3), The CA3 (x400) is characterized large and densely packed pyramidal neurons (black arrows) with apicobasal projections. The control present with characteristically defined hippocampal histomorphology with granular DG and well-defined and stained CA3. The aluminium group presents with reduced cellularity with the DG and CA3. Compared to the aluminium group, the AA, NIC and AA+NIC groups presented with improved hippocampal histomorphology and increase cellularity within the DG and CA3. The general histology of the PFC and hippocampus is demonstrated here by H and E stain.

### AA+NIC improves corticohippocampal histomorphology following aluminium-induced toxicity

The cytoprotective roles of AA+NIC on aluminium-induced histopathological alteration in the PFC and hippocampus were explored in this study. The general histology of the PFC and hippocampus following treatment with aluminium for four weeks showed that aluminium caused a significant perturbation in the general histology. At low magnification, the panoramic view of the PFC is detailed, showing the cortical layers. The control group presents with typical cortical delineation, normal cellular distribution across the layer, and characteristic staining intensity. Detailed view of the external pyramidal layer reveals well-characterized, properly distributed pyramidal neuron with, large soma, well-stained nuclei and appreciable apical and basal dendrites. The control group hippocampus at a panoramic view reveals a densely packed dentate gyrus (DG) on either side of a well-characterized cornu ammonis 3 (CA3) regions. A detailed view of the CA3 reveals large pyramidal cells with typically stained nuclei and conspicuous apical and basal projections. The histomorphology of the PFC and hippocampus of animals induced with aluminium toxicity presented with appreciable deviation from the histological quality seen in the control group. at a panoramic view, the PFC and hippocampus of the aluminium group showed a reduced staining intensity and diminished cellularity. Aluminium induced a histoarchitectural aberration in PFC cortical delineation and cellular assortment. At a detailed view, the pyramidal layers of the PFC and CA3 showed poorly characterized pyramidal cells with fragmented neuropils and poorly stained nuclei. These findings suggest that aluminium treatment perturbed the histomorphology of the PFC and hippocampus.

Treatment with AA, NIC or AA+NIC showed improved the corticohippocampal histomorphology of animal induced with aluminium toxicity. Compared to the aluminium group, the AA, NIC and AA+NIC groups presented with improved corticohippocampal histomorphology and increased cellularity.

### AA+NIC attenuates aluminium-induced corticohippocampal histopathology

Histochemical demonstration of the PFC and hippocampus of rats induced with aluminium toxicity, using Congo red and Bielschowsky’s silver stain, revealed that aluminium induced amyloid deposition and neurotic plaques formation, at prodromal stages. The Congo red of the PFC in the control group present with deeply stain neuron and pink neuropils and no pathological alteration. the aluminium group presented with deep salmon colouration across the cortical layers, suggesting the presence of amyloid plaques. In the hippocampus, however, these pathological alterations were no present. This finding suggests that, in the time frame of aluminium toxicity in this study, aluminium induced amyloidogenic changes in the PFC but not in the hippocampus. Image analysis revealed that the aluminium group had a higher staining intensity as shown by the higher mean grey area in the Image J software (figure 5). The silver stain demonstration showed an increase in pyramidal neurons with argentophilic masses and dystrophic neurons in the PFC and hippocampus of the aluminium group when compared to the control group. These increase dark colourations suggest neuropil threads formation in the cortical and hippocampal neuron following aluminium toxicity. Furthermore, immunohistochemical demonstration of the two brain areas with anti-NSE reveals that aluminium induced an increase in the expression of neuron-specific enolase, as shown by the higher NSE immunopositivity of the aluminium (figure 7). NIC alone or in combination with AA significantly reduced the amyloid plaques in the PFC. This finding suggests that that nicotine is the component of the treatment regimen that ameliorated the aluminium-induced amyloid pathology. Also, AA+NIC treatment reduced neuritic plaque formation which may be attributed to the AA therapeutic advantages. Corticohippocampal expression of NSE was also normalized following treatment with AA+NIC. These findings suggest that AA+NIC treatment synergistically ameliorated aluminium-induced corticohippocampal histopathology.

**Fig 5.**
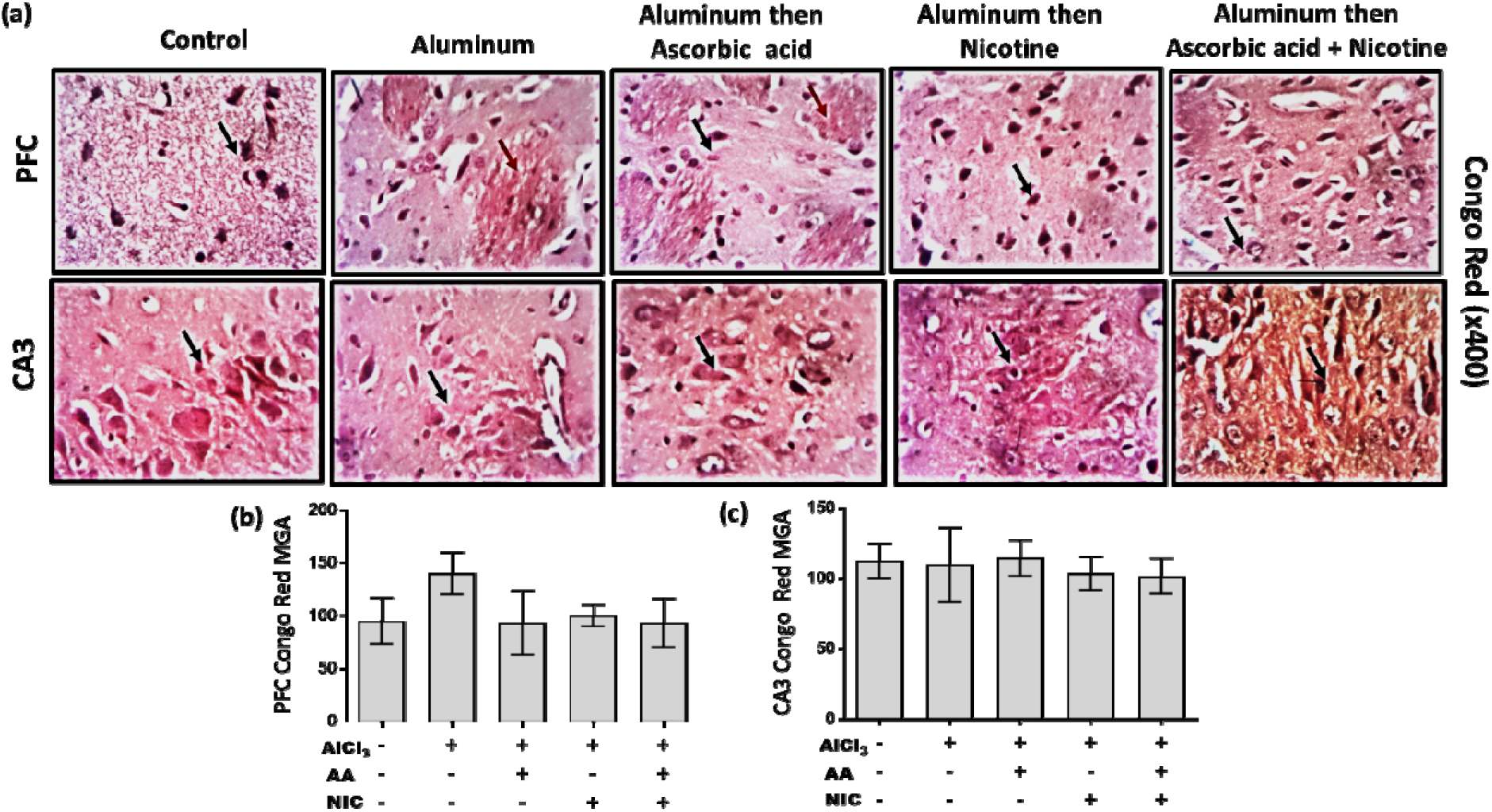
Representative photomicrographs of the amyloid (Congo red) stain (a) and respective mean grey area (staining intensity) in the PFC (b) and hippocampus (c) of Wistar rats induced with aluminium toxicity and then treated ascorbic acid, nicotine, or a combination of both. a) The PFC of the control group present with a typical distribution of pyramidal cells black arrows) in the PFC with an even pale pink background. The aluminium group present with a large salmon congophilic colouration depicting amyloidogenic deposition. The congophilic salmon colouration is reducing in the NIC and AA group and completely absent in the AA+NIC group. There are no apparent amyloid depositions in the hippocampus. The aluminium group present with the highest mean grey area in the PFC (b) and hippocampus (c) with no significant difference when compared with other experimental groups. Amyloid deposition was demonstrated by Congo red stain at x400 magnification.

**Fig 6.**
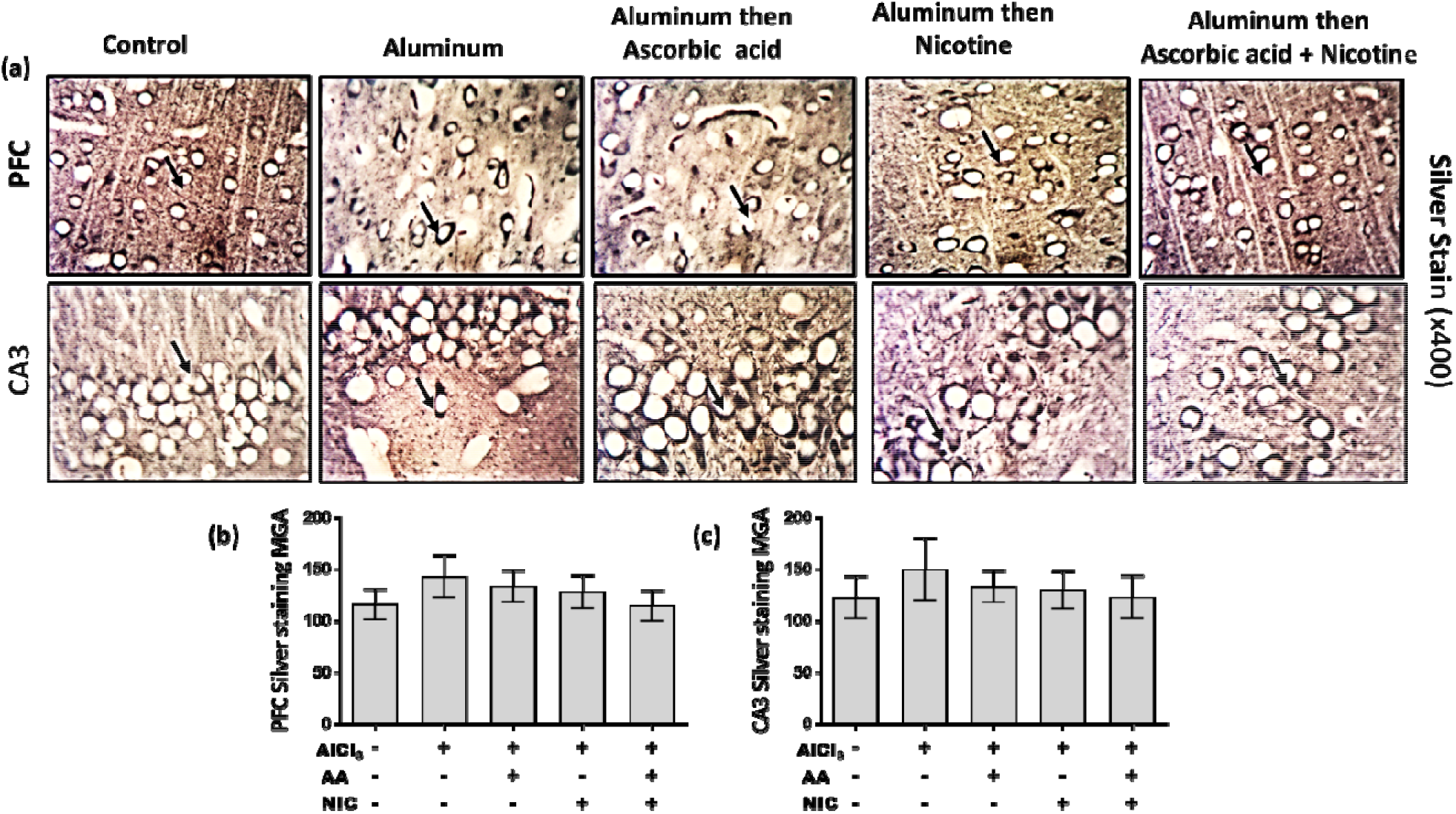
Representative photomicrographs of the silver stain (a) and respective mean grey area (staining intensity) in the PFC (b) and hippocampus (c) of Wistar rats induced with aluminium toxicity and then treated ascorbic acid, nicotine, or a combination of both. The control group present with a typically characterized distribution of pyramidal cells (black arrow) in both brain region. The cell bodies appear white with dark brown backgrounds. The aluminium group is characterized by increase argentophilic masses, neuritic threads, axonal spheroids, darkly stained neuropil around neuronal soma, suggesting tauopathic changes, similar to what is obtainable in the NIC and AA groups. This pathological alteration appears reduced in the AA+NIC group. the mean grey area in the PFC (b) and CA3 (c) was highest in the hippocampus when compared to other experimental groups. Tauopathy was demonstrated with Bielchowsky stain at x400 magnification.

**Fig 7.**
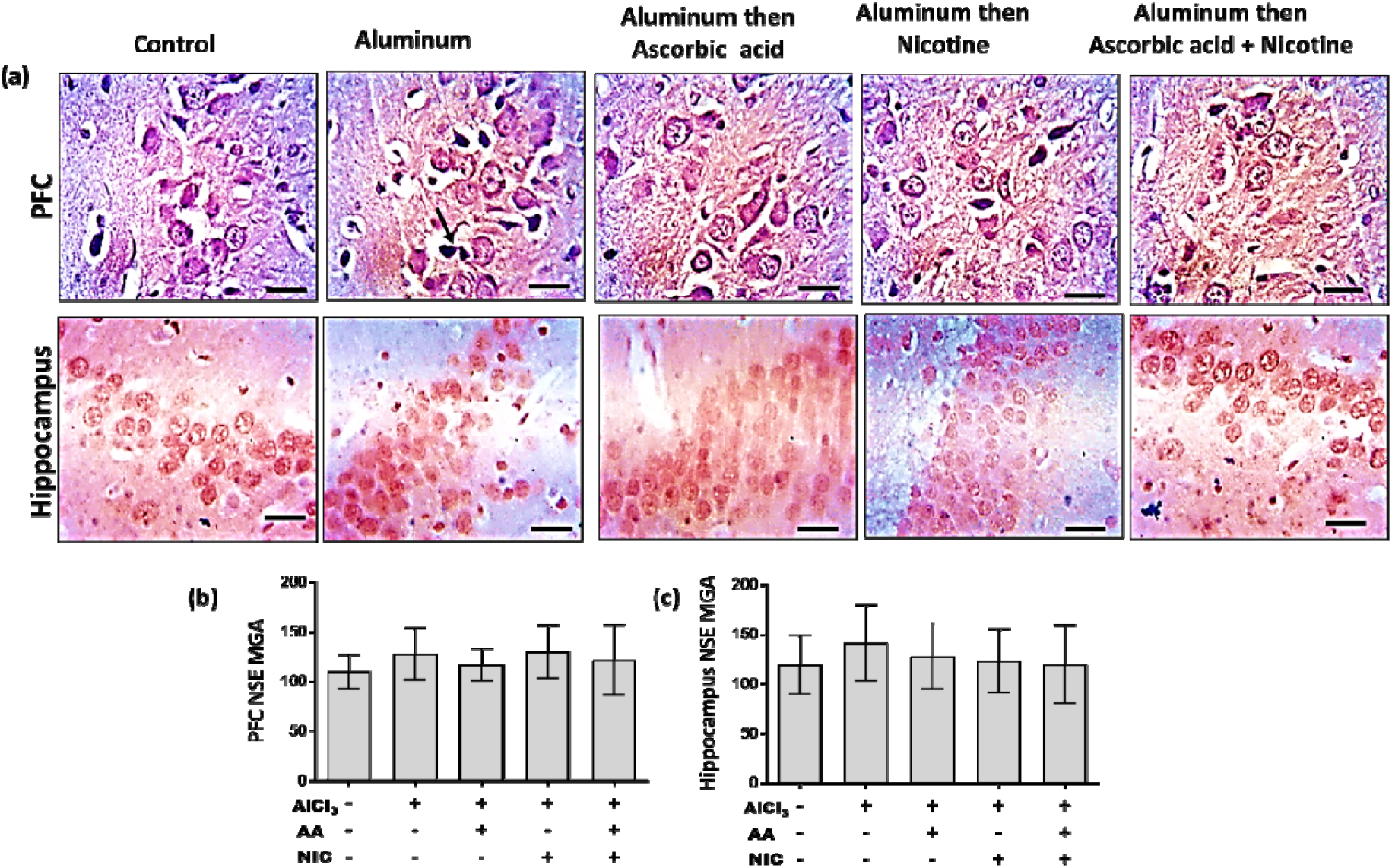
Representative photomicrography of the neuron-specific enolase immunohistochemistry (a) and respective mean grey area (staining intensity) in the PFC (b) and hippocampus (c) of Wistar rats induced with aluminium toxicity and then treated ascorbic acid, nicotine, or a combination of both. NSE immunopositivity was increased in the aluminium group compared to the control group. The AA, NIC and AA+NIC groups present normalized immunopositivity for NSE. Scale bar measures 25μ.

**Supplementary fig 1.**
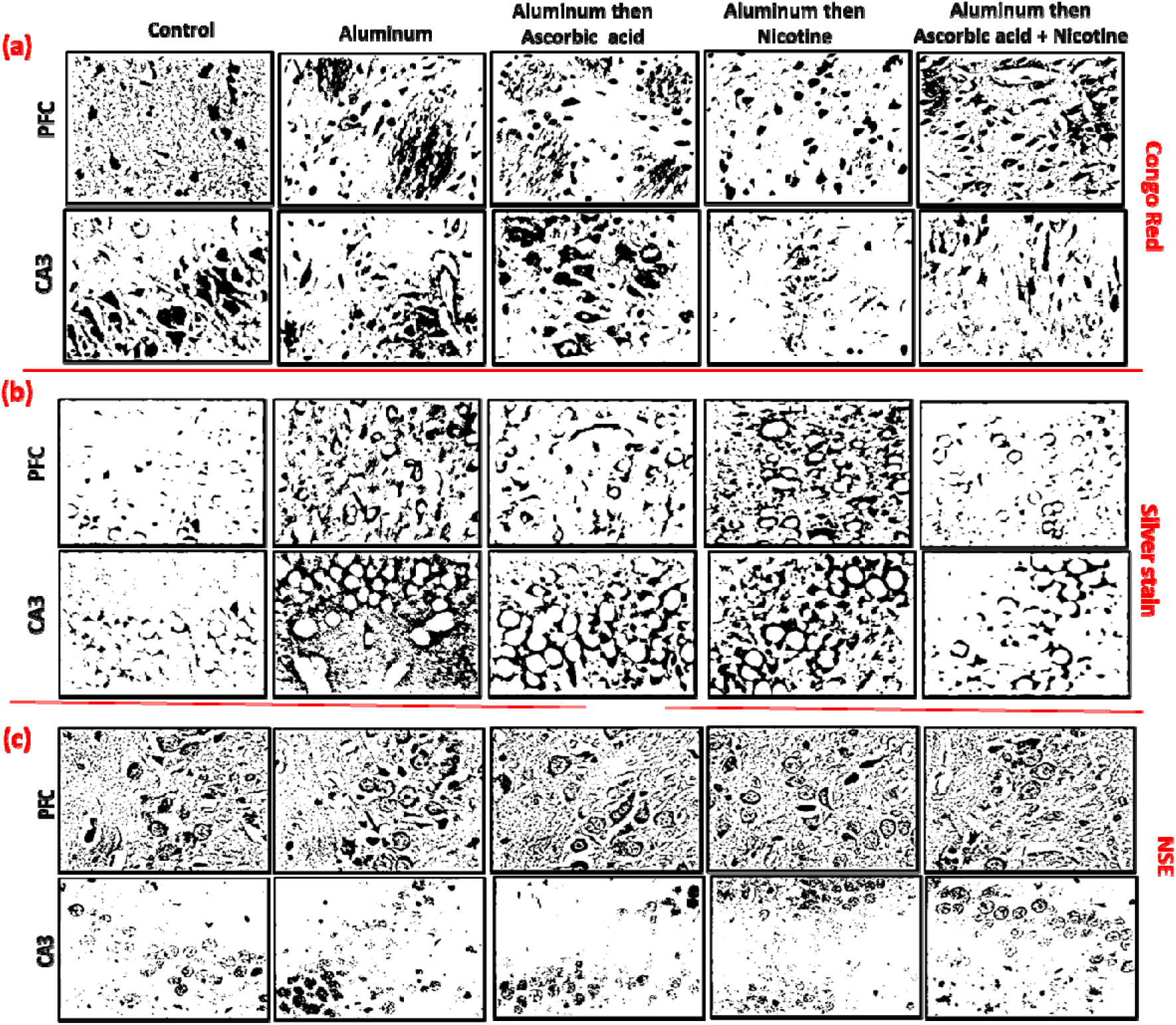
Congo red, silver stain and NSE Representative grey-scale images of used to estimate mean grey area (MGA) in image J software.

## Discussion

In the present study, by assessing body weight changes, anxiety, glucose metabolism and histopathological changes in the PFC and hippocampus of Wistar rats, we explored the therapeutic benefits of combining NIC and ascorbic acid as a treatment regimen in aluminium-induced toxicity.

Findings from this study showed that aluminium caused a significant reduction in weight gain. This finding is consistent with other reports from literature stating that chronic and acute exposure to aluminium in different kinds of experimental animals causes weight loss (Akinola et al., 2015; Elizabeth et al., 2020; Yang et al., 2017). Subchronic aluminium toxicity has been shown to cause a weight-loss by differentially altering the liver function (Azzaoui et al., 2008). Aluminium has been reported to accumulate in different organs in the body, thereby exacerbating cascades of chemical events that chelates other metals, leading to organ toxicity and ultimately reducing the functional austerity of the affected organ (Barkur and Bairy, 2015; Igbokwe et al., 2020b; Ward et al., 2001). Other studies have shown that aluminium toxicity induces a decrease in serum triglycerides and hepatic mitochondrial energy metabolism by causing dilatation of liver sinusoids, giving room for cholestasis, resulting to severely altered liver physiology and weight loss (Ibrahim et al., 2018; Kutlubay et al., 2013). In the present study, treatment of rats with AA+NIC following aluminium toxicity increased body weight. It was also observed that treatment with AA alone had a better weight enhancing effect relative to when combined with NIC. Consistent with findings from this the present study, vitamin supplementation, following aluminium toxicity restored hepatic functional integrity, which mediated an enhanced body weight gain (Kutlubay et al., 2013). Another study pointed out that dietary supplementation with AA improves the growth and general physical outcome of animals with compromised physiological wellbeing (Njoku, 1986). Microarray analysis of rat subcutaneous fat, following treatment with AA, showed that the expression of genes implicated in transcription regulation, steroidogenesis and lipid metabolism were upregulated (Campión et al., 2008). The weight enhancing effect of AA+NIC can also be attributed to the capacity of AA to effectively modulate the basal metabolic rate (Owu et al., 2006; Rucker and Steinberg, 2002)

The behavioural manifestation of experimental animals in the OFT revealed anxiogenicity of aluminium, a finding derived from the increased stretch attend posture frequency of the animal group that was treated with aluminium only. Rodents naturally elicit this risk assessment behaviour when in a new environment by performing a forward elongation of the head followed by retraction to the initial position. However, a high frequency of this behaviour shows a high level of anxiety (Holly et al., 2016). In line with findings from this study, aluminium has been reported to induce anxiety and memory decline (Crépeaux et al., 2017; Morris et al., 2017; Olajide et al., 2018). The anxiogenic propensity of aluminium can be attributed to the role it plays in mediating neurochemical disturbances and perturbation of neurotransmission in the brain (Caito and Aschner, 2015; Maya et al., 2016; Olajide et al., 2017a; Yokel, 2000). AA and NIC reduced the anxiety levels, as seen in the reduced frequency of stretch attend posture. The reduction in the stretch attend posture frequency was significant in the experimental group that was treated with both NIC and AA, relative to the aluminium group. NIC has been reported to reduced anxiety levels and improves memory function in animals stress models through modulation of neural cholinergic transmission (Costall et al., 1989; Hsu et al., 2007). Findings from human and animals studies have shown that AA has anxiolytic efficacies (de Oliveira et al., 2015; Mazloom et al., 2013). This finding is an indication of therapeutic synergy between NIC and AA in improving the behavioural outcome of the animals induced with aluminium toxicity.

In the present study, we also examined the histomorphological changes in the PFC and hippocampus of the experimental animals. In the general histology shown by H&E, the aluminium group’s PFC and hippocampus presented with reduced cellularity, distorted histoarchitectural delineation, and decreased staining intensity when panoramically viewed. A closer look at the external pyramidal layer of the PFC and pyramidal layer of the CA3 showed pyramidal cells with fragmented neuropil, vacuolated milieu, shrunken soma, inconspicuous apicobasal projections and poorly stained nuclei, all of which suggests apoptotic changes. Histochemical examination revealed congophilic plaque-like deposition in the PFC but not the hippocampus. while both brain regions were characterized by pyramidal soma with argentophilic masses, neuropil threads and dystrophic neurites in the PFC external pyramidal layer and CA3 pyramidal layer. Previous studies have also reported that aluminium toxicity induces corticohippocampal histopathologies such as chromatolysis (Olajide et al., 2017b), gliosis (Elizabeth et al., 2020), extensive neuronal cell loss (Akinola et al., 2015), neurofibrillary tangles (Kowall et al., 1989), and amyloid plaques (Rodella et al., 2008). Cumulatively, these findings have shown that aluminium induces Alzheimer-like histopathological changes in the brain of the experimental animals. AA+NIC treatment, as used in the present study, significantly improved the corticohippocampal histomorphological integrity. The experimental group post-treated with a combination of NIC and AA presented with typical general histology and characteristic corticohippocampal histoarchitecture of the PFC external pyramidal layer and CA3 pyramidal layer. The congophilic plaque-like deposition in the PFC was still present in the AA group but diminished in the NIC and AA+NIC groups, suggesting the role of NIC in decomposing amyloid plaque. NIC has been shown to amyloidosis in experimental models of AD (Lahiri et al., 2002; Nordberg et al., 2002; Utsuki et al., 2002). AA potentiates its neuroprotection, primarily by acting as an antioxidant in its capacity (Covarrubias-Pinto et al., 2015; Foyer, 2017) enabling it to scavenge for reactive oxygen and nitrogen species (Du et al., 2012), reducing lipid peroxidation (Maggio et al., 2017), and ultimately sustaining the cellular integrity of the PFC and hippocampus (Olajide et al., 2017b).

Aluminium-induced corticohippocampal histopathology can be attributed to the roles aluminium plays in exacerbating neural levels of reactive oxygen species which culminate in oxidative stress and neuroinflammation (Chakrabarty et al., 2012; Mustafa Rizvi et al., 2014; Rodella et al., 2008). Aluminium has been reported to induce excess iron uptake in neurons (Greger et al., 1997) which results in distortion of the intracellular labile iron pool (Breuer et al., 2008), and then drive cascade of chemical events that accelerate ROS production, beyond what endogenous antioxidant enzymes can cope with (Igbokwe et al., 2020b). Aluminium-induced excess oxidative stress culminates in neurotoxic events, such as lipid peroxidation (Bhattacharjee et al., 2014), neuroinflammation (Yegambaram et al., 2015) and glia activation (Akinrinade et al., 2015), that ultimately compromises structural and functional integrity of neural cells, hence the manifestation of the behavioural decline seen in the animals exposed to aluminium toxicity. All the aforementioned chemical events following aluminium toxicity, especially neuroinflammation, require a huge amount of cellular energy to be executed (Corrado et al., 2012). There is a documented molecular interplay between oxidative stress and glucose bioenergetics (Balazs and Leon, 1994; Bigl et al., 1999). This is evidenced in this study by the depleted activities of G6PDH and elevated lactate activities in the PFC and hippocampus following oral infusion of aluminium. This finding implies that aluminium may have induced an energy metabolic reprogramming in the studied brain areas, from oxidative phosphorylation to glycolysis, a phenomenon known as the Warburg effect (Otto, 2016). The speculated metabolic paradigm shift often occurs in a bid to cater for the excess energy demand in response to neuroinflammation (Missiroli et al., 2020). G6PDH is a rate-limiting enzyme that catalyzes the irreversible conversion of glucose-6-phosphate to 6-phosphogluconolactone in pentose phosphate pathway (Ulusu, 2015; Zhang et al., 2014). Due to the essentiality of the G6PDH in the glycolytic pathway, it’s deficiency has been reported to promote oxidative stress, because the reduced nicotinamide adenine dinucleotide phosphate (NADPH) maintained by this enzyme is important in supplying glutathione to cells, for mopping up of excess highly reactive hydroxyl ions (Mejías et al., 2006). In fact, G6PDH deficiency and AD have colloquially been regarded as “partners in crime” (Ulusu, 2015).

Aluminium-induced activation of glial cells (Elizabeth et al., 2020) causes conformational changes in neuroenergetics during neuroinflammation by shunting more pyruvate for glycolytic conversion to lactate (Mason, 2017), thereby availing more ATP to reactive astrocyte. Despite the advantage of the metabolic reprogramming, sustained lactate accumulation has been documented to exacerbate amyloidogenesis by promoting molecular interaction between endoplasmic reticulum chaperone and APP (Xiang et al., 2010), which may explain the increase argentophilic bodies around the soma of corticohippocampal neurons. Increase lactate activity, as induced by aluminium in this study, has also been reported to be an indicator of cellular hypoxia and promote cell death (Firth et al., 1995; Koukourakis et al., 2003; Rama Rao and Kielian, 2015), which may justify the observed perturbed corticohippocampal histomorphology. An earlier study reported a link between increased lactate dehydrogenase and improved intracellular NSE expression in neurotoxin-exposed cells (Thomas et al., 1991). The production and function of NSE in glial activation is enhanced, with major factors involved in neurodegenerative diseases (Chen et al., 2018; Zabel and Kirsch, 2013). PFC and hippocampus Immunohistochemical analysis showed an increase in the expression of the NSE, though not statistically significant when qualitatively examined. NSE, a glycolytic enzyme that can play a dual role in promoting neuroinflammation and neuroprotection in neuronal tissues (Haque et al., 2018), is expressed during morphological changes in reactive astrocytes, following activation by neuroinflammatory stimulus (Vinores and Rubinstein, 1985). Hence, we speculated that the observed increase in NSE expression may be due to the emergence of activated astrocyte following aluminium toxicity. NSE’s differential expression may be linked to astrocyte degenerative or neuroprotective roles in neuronal injury because reactive astrocytes lead to neuronal death with impaired structural integrity in neurodegenerative disorders (Sensenbrenner et al., 1997; Steiner et al., 2006; Yamamoto and Kawana, 1991).

In the present study, G-6-PDH and lactate levels were significantly normalized by the AA+NIC treatment regimen, primarily through the therapeutic benefits of AA. Coupled with the improved corticohippocampal histomorphology, NSE expression was also normalized following treatment with AA+NIC. Chronic administration of NIC, *in vivo*, in an experimental AD model diminished Aβ-induced behavioural decline by reducing BACE-1 expression and Aβ levels in addition to modulating neural nicotinic receptors (Srivareerat et al., 2011). Besides, in transgenic mice expressing NSE, differential doses treatment of NIC displayed an improvement in memory and increased the expression of nicotinic receptors (Shim et al., 2008). These findings explain how nicotine is able to elicit its therapeutic advantages. However, NIC, as part of the treatment scheme, promotes the generation of ROS (Gbadamosi et al., 2019, 2016; Omotoso et al., 2018) and potentially neurotoxic (Carlson et al., 2001; Mishra et al., 2015). Nevertheless, due to its powerful antioxidant capacity, AA can counterbalance NIC’s neurotoxic tendencies while providing a degree of quality neuroprotection against corticohippocampal toxicity induced by aluminium. In an earlier study with Olajide et al., we examined AA’s roles in behavioural and corticohippocampal neurochemical changes caused by acute aluminium toxicity in rats and found that a two-week treatment with AA expressively reduced rat behavioural deficits through its ability to scavenge free radicals, prevent excessive lipid peroxidation while modulating neuronal bioenergetics (Olajide et al., 2017b). Kazuma and colleagues had earlier reported that AA restored behavioural deficit in addition to depleting amyloid plaques in a mouse model of AD, and attributed their findings to decreases CNS oxidative damage mediated by AA (Murakami et al., 2011). AA has been reported to protect against apoptosis by regulating the modulation of the mitochondrial pathway of cytochrome C activities that resulted in the altered release of lactate dehydrogenase (Chen et al., 2018).

## Conclusion

This study has investigated the therapeutic benefits of incorporating NIC and ascorbic acid as a treatment regimen in aluminium-induced toxicity by analyzing changes in body weight, fear, glucose metabolism and histopathological changes in the PFC and hippocampus of Wistar rats. Based on the findings from this study, combining AA and NIC as a treatment regimen has more advantage in the sense that, while nicotine reduced aluminium-induced corticohippocampal congophilicity, ascorbic acid significantly improved energy metabolism, leading to enhanced corticohippocampal histomorphology and reduced anxiety. However, more studies need to be carried out to elucidate the roles of AA+NIC on activities of inflammatory cytokines, antioxidant enzymes, glial activation and synaptic morphology within the PFC and hippocampus following aluminium-induced neuroinflammation. Also, in silico modelling of a complex between nicotine and ascorbic acid will provide insights on its putative pharmacological advantages in modulating pathways implicated in AD.

## Conflict of interest

The authors declare no conflict of interest

## Notes

### Competing Interest Statement

The authors have declared no competing interest.

### Summary of Updates

The manuscript was revised for punctuation omission between aluminium and induced

